# Experimental evidence that investment into ejaculation and mating effort impose different life history trade-offs

**DOI:** 10.1101/2024.01.26.577329

**Authors:** Meng-Han Joseph Chung, Rebecca J. Fox, Michael D. Jennions

## Abstract

When males compete, sexual selection favors reproductive traits that increase their mating or fertilization success (pre- and post-copulatory sexual selection). It is assumed that males face a trade-off between these two types of sexual traits because they both draw on investment from the same pool of resources. Consequently, investment into mate acquisition or ejaculation should create similar trade-offs with other key life history traits. Tests of these assumptions are exceedingly rare. Males only ejaculate after they mate, and investment into ejaculation is therefore highly correlated with mating effort. Consequently, little is known about how each component of reproductive investment affects a male’s future performance. Here, we ran an experiment using a novel technique to distinguish the life history costs of mating effort and ejaculation for mosquitofish (*Gambusia holbrooki*). We compared manipulated males (mate without ejaculation), control males (mate and ejaculate) and naïve males (neither mate nor ejaculate) continuously housed with a female and two rival males. We assessed their growth, somatic maintenance, mating and fighting behavior, and sperm traits after 8 and 16 weeks. Past mating effort significantly lowered a male’s future mating effort and growth, but not his sperm production; while past sperm release significantly lowered a male’s future ejaculate quantity, but not his mating effort. These findings challenge the assumption that male reproductive investment draws from a common pool of resources such that, all else being equal, greater mating effort reduces ejaculate investment, and *vice versa*. Instead, we provide clear evidence that investment in traits under pre- and post-copulatory sexual selection have different trait-specific effects on subsequent male reproductive performance that occur both early and late in life.

## Introduction

Accounting for variation in life-histories among individuals, populations or species is a central challenge in evolutionary biology [1–3]. A key to understand this variation is to recognize that individuals have limited resources to allocate to traits. Time, material or energy invested in one trait often lowers investment in another. This creates life history trade-offs such that fitness is maximized by the optimal allocation of resources. Notably, investment by males into sexually selected traits that increase mating success lowers investment into naturally selected traits or somatic maintenance [4–6]. Sexually selected traits such as extravagant ornaments or weapons, combat with rivals, or engaging in courtship often elevate mortality [7–9] because they are energetically expensive [10], or activate physiological pathways with harmful pleiotropic effects on somatic maintenance (e.g., immunosuppression; [11, 12]). Far less is known about the life history costs of traits under post-copulatory sexual selection that increase fertilization success when females mate multiply and sperm compete (e.g., larger testes, bigger ejaculates; [13, 14]). Although it is well established that ‘sperm is not cheap’ [15–17], the life-history costs of increased ejaculate investment under competition are often unclear [18].

Theoretical models of sperm competition games predict a direct trade-off between traits that enhance mate acquisition and traits that elevate sperm competitiveness based on the assumption that both types of traits draw from a single, common pool of resources [19–21]. In support of this assumption, some cross-species studies report a negative relationship between pre- and post-copulatory sexual traits after controlling for confounding factors that affect the benefits of investment (e.g., how readily females are monopolized by males; [22]). However, not all traits under pre-copulatory sexual selection seem to trade off with ejaculate traits (e.g., ornaments do, but weapons do not, in primates; [23]). Basic physiology provides an obvious reason to challenge the assumption that investment into ejaculates and mating effort is equivalent. Sperm production depends heavily on meiosis in germline tissues [24], while mate acquisition depends on investment in somatic tissues maintained by mitosis (e.g., the production of physical traits or the use of muscles in sexual displays or combat; [25, 26]). This raises the issue of whether it is appropriate to assume that ejaculates and mating effort draw from a single pool of resources. Predictions about life history variation change if ejaculates and mating effort impose different trade-offs.

To date, very few studies have used an experiment to compare the effect of pre- versus postcopulatory sexually selected traits on life history trade-offs. These trade-offs cannot be reliably detected by comparing mated and virgin individuals [27–29]: mated males who ejaculate (and then replenish sperm) have already paid the preceding costs of mate acquisition. More generally, males with greater pre-copulatory investment (i.e., larger sexual ornaments, higher courtship rate) have greater mating success, which, in turn, results in more frequent sperm replenishment [30]. Mating effort and ejaculation are therefore highly correlated and difficult to separate statistically. Ultimately, an experimental approach is essential to determine the causal effects of mating effort and ejaculation on each other, and on other traits. However, such experiments are lacking due to the logistic challenge of manipulating the ejaculate rate without changing the mating rate.

We have developed a surgical technique for mosquitofish, *Gambusia holbrooki*, that provides a way to address the question of the relative life history costs of ejaculation and mating effort [31]: we prevent males from ejaculating by ablating the tip of their intromittent organ. Ablated males behave normally and try to mate but they do not receive the sensory cues that induce sperm release [31]. Surgery by itself does not affect male attractiveness [32], male-male aggression (this study) or baseline sperm production [33]. Male *G. holbrooki* attempt to mate every few minutes [34]; and females mate multiply [35], which selects for large ejaculates [14]. Ejaculate production seems costly as greater food availability increases ejaculate size and hastens sperm replenishment [36]. Strong competition for mates and high sperm competition make *G. holbrooki* an ideal species to study the costs of male investment into sexually selected traits.

We randomly assigned males to three treatments to independently manipulate mating effort and the frequency of sperm release/replenishment. To quantify the cost of mating effort, we compared (a) *naïve* males without access to females and rivals that neither attempted to mate nor ejaculated, to (b) *ablated* males with access to females and male rivals that could mate/fight but not ejaculate. To quantify the cost of sperm release/replenishment, we compared (b) *ablated* males to (c) *non-ablated* males with access to females and rivals that could both mate/fight *and* ejaculate. To quantify trade-offs, we measured both somatic traits (growth, immune response) and reproductive traits (mating and aggressive behavior, sperm traits) after 8 and 16 weeks. Mating and fighting behavior were measured under standardized conditions: the focal male was placed in a tank containing a new female and a novel rival male for behavioral observation. Ejaculates were also measured under standardized conditions: to control for treatment-induced variation in a male’s recent history of mating and ejaculation, the focal male was stripped of sperm and isolated for 5 days [36] before we stripped his sperm again to quantify the accumulated sperm reserves (*total sperm count* and *sperm velocity*); after 24 hours, we again stripped sperm from the male to measure his *rate of sperm replenishment*.

If mating effort and ejaculation draw on the same (or a highly overlapping) pool of resources, all else being equal, we predict that:

1. Increased mating effort will lower future sperm production, and *vice versa*.
2. Mating effort and ejaculation will affect the same naturally selected traits.
3. The costs of mating effort and ejaculation will cumulatively increase and be more apparent in older males.

Our current study extends an earlier study [32] in two key ways. First, all focal males were housed with a female and two rival males to provide social cues about sexual competition. In the earlier study, focal males lacked rivals and social cues about sexual competition. Their absence might have lowered reproductive investment, making it harder to detect life history trade-offs. Second, we measured costs at 16 weeks. Previously we only measured costs after 8 weeks [32]. Adult males can live up to 20 weeks in the wild [37]. In our current study, we can therefore test for costs of reproductive investment that might be absent in younger males. In general, the costs of reproduction accelerate with age (i.e. senescence occurs) [3, 38].

## Results

### Preliminary test: no surgery effect on performance during male-male interactions

To test whether the ablation surgery itself causes behavioral changes in the current experiment where males compete with rivals, we haphazardly assigned virgin males to be either *ablated* or *non-ablated* (two-sample *t*-test for body size: *t* = 0.517, p = .606; *n* = 70 per group). We measured mating behavior before and after each male was housed with a female and two rival males for seven days of sexual interactions. Previous studies have shown that seven days suffices to detect behavioral adjustments in response to past sperm investment [39] or past mating effort [40].

There was no detectable effect of ablation surgery on male behavior (Table 1), except that *ablated* males made fewer mating attempts on Day 0. However, this effect was undetectable after seven days (Table 1), indicating no long-term effects of surgery.

**Table 1.** Surgery effect (i.e., gonopodium ablated or intact) on mating behaviors. Given the significant interaction between gonopodium state and day on *number of mating attempts*, the surgery effect was tested separately on each day. Statistical outputs of final models excluding non-significant interactions are shown. Full models are provided in S1 Table. The bold font highlights significance at the 0.05 level.

| Trait | Predictor | Test statistic | p |
| --- | --- | --- | --- |
| **Number of nips and approaches** | Body length * Day | $\chi^2_1 = 4.248$ | **.039** |
| | Gonopodium state | $\chi^2_1 = 0.282$ | .596 |
| | Day | $\chi^2_1 = 47.419$ | **< .001** |
| | Body length (standardized) | $\chi^2_1 = 2.965$ | .085 |
| | Male size difference (standardized) | $\chi^2_1 = 7.625$ | **.006** |
| **Time spent near female** | Gonopodium state | $F_{1,139.06} = 0.220$ | .639 |
| | Day | $F_{1,134.21} = 0.767$ | .383 |
| | Body length (standardized) | $F_{1,188.74} = 5.010$ | **.026** |
| | Male size difference (standardized) | $F_{1,238.52} = 0.332$ | .565 |
| **Number of mating attempts** | Gonopodium state * Day | $\chi^2_1 = 5.838$ | **.016** |
| | Gonopodium state | $\chi^2_1 = 2.372$ | .124 |
| | Day | $\chi^2_1 = 33.468$ | **< .001** |
| | Body length (standardized) | $\chi^2_1 < 0.001$ | .996 |
| | Male size difference (standardized) | $\chi^2_1 = 1.460$ | .227 |
| **First day of sexual interaction (Day 0)** | Gonopodium state * Body length | $\chi^2_1 = 0.028$ | .867 |
| | Gonopodium state | $\chi^2_1 = 5.030$ | **.025** |
| | Body length (standardized) | $\chi^2_1 = 0.135$ | .713 |
| | Male size difference (standardized) | $\chi^2_1 = 0.741$ | .389 |
| **Seven days after sexual interaction (Day 7)** | Gonopodium state * Body length | $\chi^2_1 = 0.093$ | .761 |
| | Gonopodium state | $\chi^2_1 = 1.812$ | .178 |
| | Body length (standardized) | $\chi^2_1 = 0.238$ | .625 |
| | Male size difference (standardized) | $\chi^2_1 = 3.428$ | .064 |

### The effect of past reproductive investment on somatic traits

Controlling for initial body size, males who had invested in mating effort (i.e., *ablated* or *non-ablated* males) were significantly shorter and thinner at week 8 (Fig 1A and 1B), measured as either standard length (SL: snout tip to base of caudal fin) or body depth (BD: base of dorsal fin to ventral side of body) (Tukey’s tests: vs *naïve* males; all p < .001; S2 Table). In contrast, whether a male released sperm in the first 8 weeks did not affect his SL or BD (Tukey’s tests: *ablated* vs *non-ablated* males, both p > 0.25). Males barely grew from weeks 8 to 16 (Fig 1); hence, there was no effect of past reproductive investment on body size (S2 Table).

**Fig 1.**
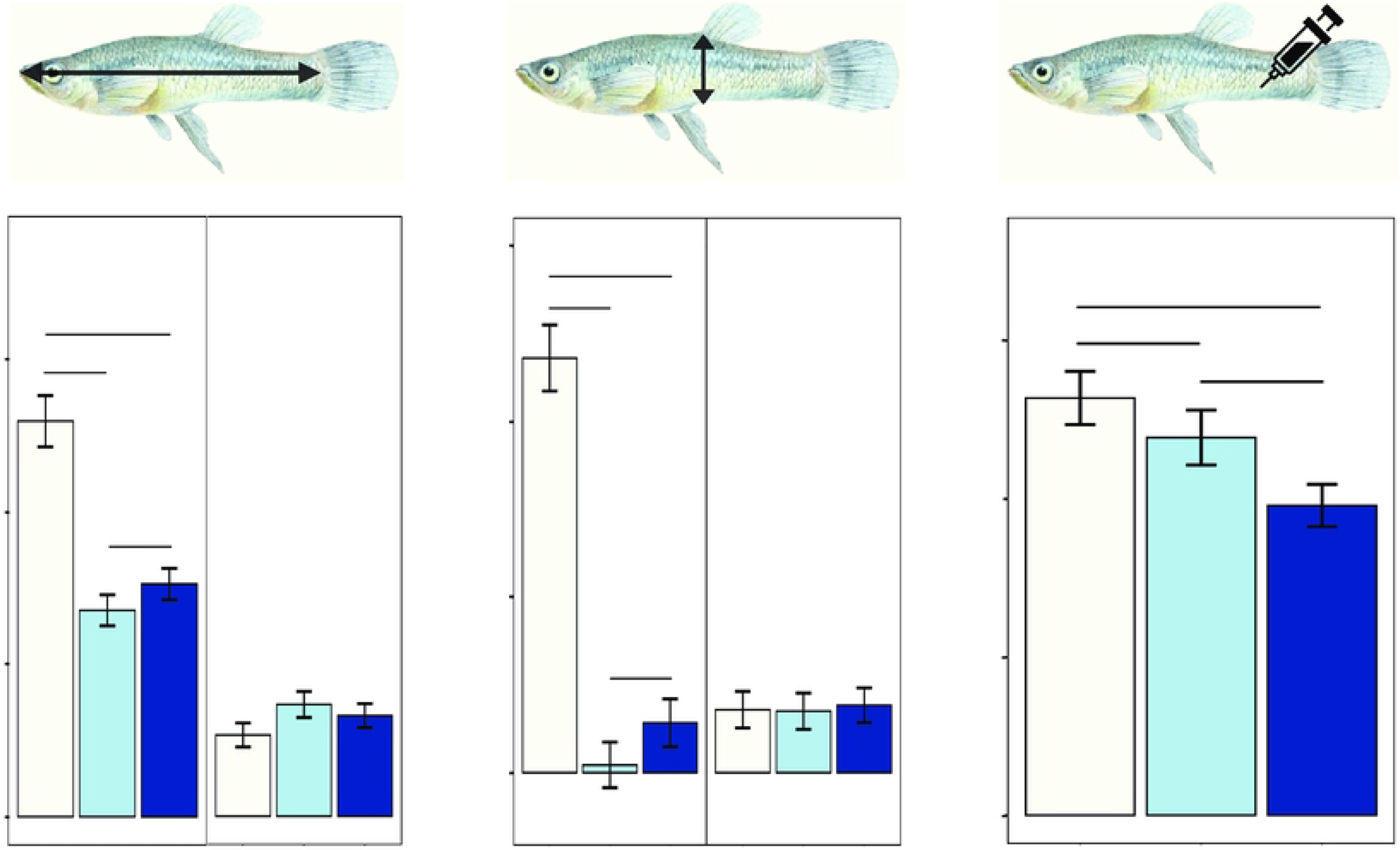
Effects of reproductive treatment and age at testing on somatic traits. (A) standard length, (B) body depth and (C) immune response. When there is a significant treatment*age interaction (see S2 Table for details), data at weeks 8 and 16 are presented separately. Light blue = *naïve* males that neither mated nor ejaculated; blue = *ablated* males that mated but did not ejaculate (hence low/no sperm replenishment); dark blue = *non-ablated* males that both mated and ejaculated. Significant level: *** p < .001; ** p = .001 to .01; * p = .01 to .05; n.s. = non-significant. P-values are from Tukey’s tests (see text). Data is shown as mean ± SE.

Past reproductive investment significantly affected immune response (S2 Table). *Naïve* males had greater immunocompetence than *non-ablated* males, who had invested into both mating effort and ejaculation (Tukey’s test: p = .013; Fig 1C). However, neither greater mating effort alone (*naïve* versus *ablated* males) nor frequent sperm release alone (*ablated* versus *non-ablated* males) lowered immunocompetence (Tukey’s tests: both p > .165). These treatment effects did not differ between weeks 8 and 16 (interaction: p = .538), nor did immune function decline with age (p = .401) (S2 Table).

### The effect of past reproductive investment on reproductive traits

We tested how past reproductive investment affected a male’s subsequent mating performance by observing his behavior towards a female in the presence of a novel rival after 8 and 16 weeks. Past reproductive investment and age at testing did not interact to affect any of the reproductive traits (S3 Table), but both factors independently affected male behavior (Table 2). *Naïve* males approached and nipped a rival significantly more often than males with greater past mating effort (i.e., *ablated* or *non-ablated*) (Tukey’s tests: both p < .001; Fig 2A), but repeated sperm release in the past did not affect male aggression (Tukey’s test: *ablated* vs *non-ablated* males, p = .738). Similarly, *naïve* males spent significantly more time near females than did males with higher past mating effort (Tukey’s tests: both p < .001; Fig 2B, Table 2), but past sperm release did not affect the time spent near females (Tukey’s test: p = .061). Whether or not a male attempted to mate was unaffected by his past reproductive investment (Table 2). Of those males that attempted to mate, males with greater past mating effort made significantly fewer attempts (*ablated* or *non-ablated* vs *naïve* males, Tukey’s tests: both p < .001; Fig 2C). In contrast, past sperm release did not affect the number of mating attempts (Tukey’s test: p = .732). Age at testing had no effect on the number of mating attempts or time spent near females, but older males were significantly more aggressive towards rivals (weeks 8 vs 16, Table 2).

**Fig 2.**
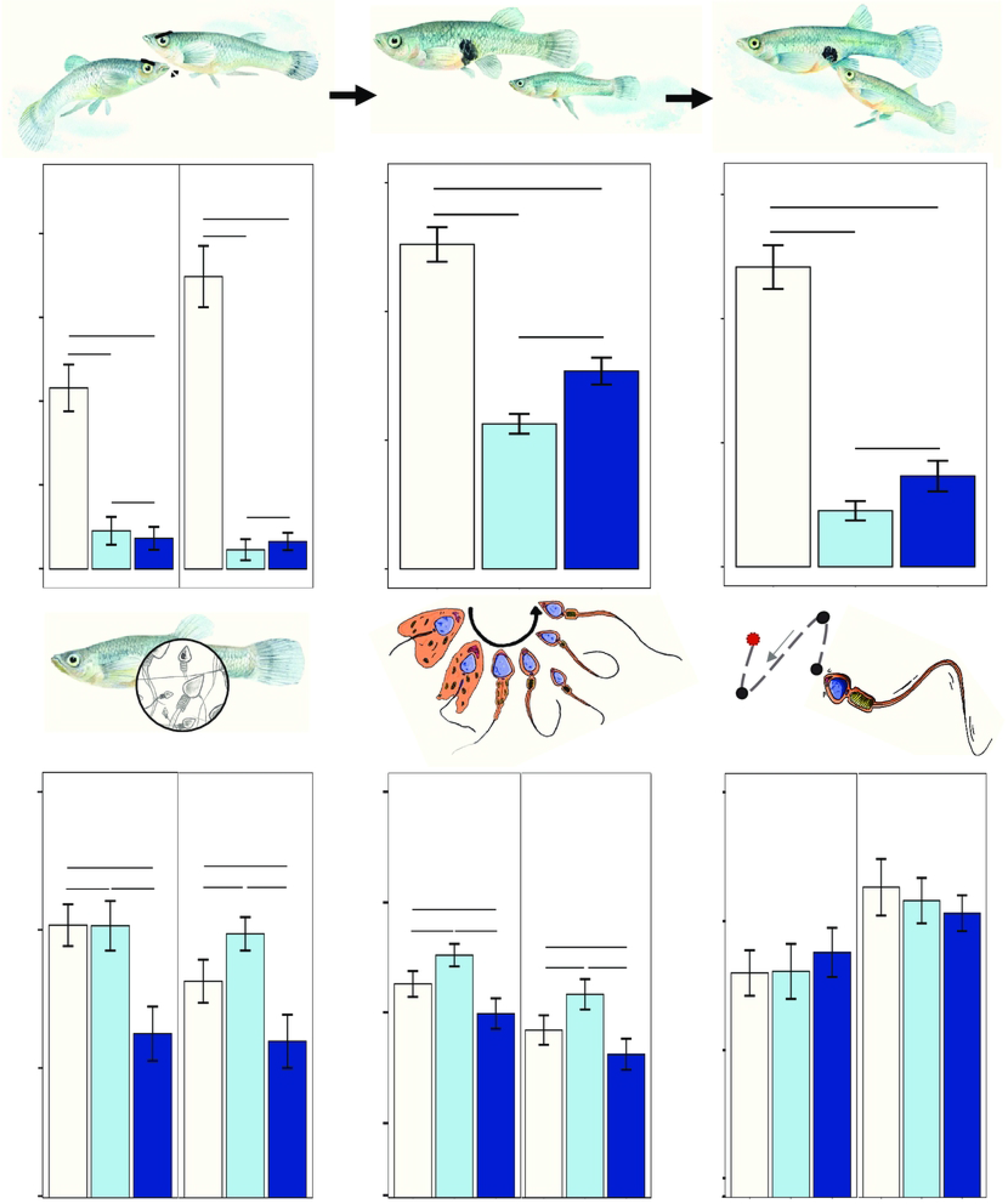
Effects of reproductive treatment and age at testing on reproductive traits. Mating performance: (A) number of rapid approach and nips to rival, (B) time spent near female, (C) number of mating attempts; and ejaculate traits: (D) total sperm number (log-transformed), (E) number of replenished sperm (log-transformed), (F) sperm velocity. Given a significant effect of age (see Table 2 for details), data at weeks 8 and 16 are presented separately. Light blue = *naïve* males; blue = *ablated* males; dark blue = *non-ablated* males. Significant level: *** p < .001; ** p = .001 to .01; * p = .01 to .05; n.s. = non-significant. P-values are from Tukey’s tests (see text). Data is shown as mean ± SE.

**Table 2.** Effects of reproductive treatment and age at testing on male reproductive traits. Main effects are obtained from final models excluding any non-significant interactions. Statistical outputs of initial models with the interaction are provided in S3 Table.

| Trait | Predictor | Test statistic | p |
| --- | --- | --- | --- |
| ***Mating performance*** |  |  |  |
| **No. approaches and nips** | Reproductive treatment | $\chi^2_2 = 192.298$ | < **.001** |
| | Age at testing | $\chi^2_1 = 5.533$ | **.019** |
| | Male size difference (standardized) | $\chi^2_1 = 0.668$ | .414 |
| | Initial body length (standardized) | $\chi^2_1 = 3.150$ | .076 |
| **Time spent near female** | Reproductive treatment | $F_{2,156.16} = 40.033$ | < **.001** |
| | Age at testing | $F_{1,173.29} = 0.008$ | .929 |
| | Male size difference (standardized) | $F_{1,302.82} = 0.446$ | .505 |
| | Initial body length (standardized) | $F_{1,231.28} = 1.725$ | .190 |
| **No. mating attempts** |  |  |  |
| Zero-inflation | Reproductive treatment | $\chi^2_2 = 4.916$ | .086 |
| | Age at testing | $\chi^2_1 = 3.714$ | .054 |
| | Male size difference (standardized) | $\chi^2_1 = 0.007$ | .932 |
| | Initial body length (standardized) | $\chi^2_1 = 1.158$ | .282 |
| Conditional | Reproductive treatment | $\chi^2_2 = 33.512$ | < **.001** |
| | Age at testing | $\chi^2_1 = 0.596$ | .440 |
| | Male size difference (standardized) | $\chi^2_1 = 3.657$ | .056 |
| | Initial body length (standardized) | $\chi^2_1 = 6.614$ | **.010** |
| ***Ejaculate traits*** |  |  |  |
| **Total sperm count** | Reproductive treatment | $F_{2,152.28} = 13.731$ | < **.001** |
| | Age at testing | $F_{1,176.66} = 8.587$ | **.004** |
| | Initial body length (standardized) | $F_{1,169.44} = 59.365$ | < **.001** |
| **Sperm replenishment rate** | Reproductive treatment | $F_{2,149.77} = 10.155$ | < **.001** |
| | Age at testing | $F_{1,169.65} = 35.127$ | < **.001** |
| | Initial body length (standardized) | $F_{1,164.76} = 37.206$ | < **.001** |
| **Sperm velocity (VCL)** | Reproductive treatment | $F_{2,295} = 0.061$ | .941 |
| | Age at testing | $F_{1,295} = 6.512$ | **.011** |
| | Initial body length (standardized) | $F_{1,295} = 13.370$ | < **.001** |

Ejaculate quantity (total sperm count and sperm replenishment rate), but not sperm velocity, was significantly affected by past reproductive investment (Table 2). In contrast to mating behavior, however, the decline in total sperm count and sperm replenishment rate were due to past sperm release (Tukey’s test: *ablated* vs *non-ablated*, both p < .001; Fig 2D and 2E), but not past mating effort (Tukey’s test: *naïve* versus *ablated,* p = .661 and .082). Finally, older males had a significantly lower total sperm count and a slower rate of sperm replenishment, but they produced sperm that swam significantly faster (Table 2).

## Discussion

Current reproductive investment is assumed to impose costs by depleting a common pool of resources that are also used for growth, maintenance and future reproduction [41–43]. This creates life history trade-offs among different components of fitness. Similarly, there are trade-offs among traits, including ones that affect the same fitness component. Specifically, for sexually selected traits that increase the rate of reproduction, ‘sperm competition game’ models of optimal investment make the assumption that “*each male has a fixed total energy budget for reproduction (R) which he divides between his allocation to precopulatory competition (T) and his allocation to postcopulatory competition (U), so that: R = T + U*” [20, 21]. Hence, greater mating effort should reduce ejaculate production, and *vice versa* [44, 45]. There is seemingly empirical support for this prediction. For example, male houbara bustards with a higher rate of sexual displays had a more rapid decline in ejaculate viability with age [46]. However, these males would have ejaculated more frequently because females prefer males with greater courtship effort [47]. In observational studies, distinguishing how pre- and post-copulatory reproductive investment affects life history trade-offs is therefore challenging because investment in mate acquisition always precedes ejaculation.

By experimentally manipulating pre- and post-copulatory investment in *G. holbrooki*, our results reveal that past sperm release only affects future ejaculate production, while past mating effort only affects future mating performance and growth. We can therefore reject our first two predictions that: (1) greater mating effort will lower future sperm production, and *vice versa*; and that (2) mating effort and ejaculation will affect the same naturally selected traits. Our findings offer insights into how to apply life-history trade-offs to different sexually selected traits, and illuminate hypotheses about differences among traits in their age-related decline in performance (i.e., senescence) [48, 49]. For example, greater allocation to ejaculate size due to a higher perceived risk of sperm competition [13] could accelerate the senescence of ejaculate quality, but have no impact on the courtship or fighting ability of older males.

### Mating effort and sperm release have different effects on later-life performance

Why did past mating effort reduce male growth? First, spending more time pursuing females and attacking rivals reduces the time available to forage. In guppies, males spent 65-75% less time foraging when trying to mate [50], and male mosquitofish are likely to experience a similar time trade-off. Second, the expression of sexual behaviors often elevates testosterone levels [51, 52], which tends to be associated with a higher metabolic rate and energy consumption [53]. Even so, it is difficult to explain the effect of mating effort on growth solely by invoking energy availability as past mating effort did not affect future sperm production, which also has energetic costs [17]. Another explanation why mating effort reduced growth is that prolonged locomotion when chasing females generated reactive oxygen species, which damaged somatic cells and muscles [54]. Damage to somatic tissue could also explain the decline in future mating performance, but not in future sperm production if germline tissue remains unaffected. Another possibility is that ablated males strategically, and plastically, decrease their mating effort because they detected their failure to inseminate females, possibly based on the lack of decline in their sperm reserves. This seems unlikely, however, given our preliminary test showed no short-term decline in mating effort by ablated males that did not ejaculate (Table 1).

Similarly, why did sperm release and replenishment affect future ejaculate production, but not future mating or fighting behavior, nor growth? Following the logic outlined above, this might reflect costly damage that is localized to germline tissue, such as toxic metabolic waste products accumulating in the testes [24], but has no effect on somatic tissue. Our finding that past investment in ejaculation lowered future sperm quantity, but not sperm velocity, suggests that sperm quantity is more strongly condition-dependent [55]. One possibility is that males benefit from an allocation strategy that adaptively shifts resources to maintain sperm quality, even if this lowers their sperm count. In support of this explanation, we have shown that, controlling for sperm count, the sperm of mated and virgin male mosquitofish is equally effective for artificial insemination [56], suggesting no difference in sperm quality.

### Age and reproductive decline

Given the importance of reproduction in determining lifetime fitness, one might expect little reproductive decline earlier in adulthood, but a steep decline as males approach their maximum natural lifespan [57, 58]. However, we detected a significant phenotypic decline in most reproductive traits after only 8 weeks of reproductive investment (Fig 2), indicating that costs arise shortly after males start to engage in reproduction. There were no significant interactions between the reproductive treatment (naive, ablated, non-ablated) and age (weeks 8 or 16) for any of the measured reproductive traits (Table 2), indicating that the main effects of mating effort and ejaculation did not increase for older males. We can therefore reject our third prediction that the costs of mating effort and ejaculation will cumulatively increase and be more apparent in older males. Even so, the absolute sperm count and rate of sperm replenishment was generally lower in older males (Fig 2). Conversely, however, older males had faster swimming sperm and were more aggressive to rivals. There are therefore effects of male age itself (or correlates thereof) which are independent of the cumulative effects of mating effort and ejaculation. Our findings indicate that male *G. holbrooki* pursue a life history strategy with the potential for high reproductive success early in life, even if this creates intensive wear-and-tear [5, 46, 59, 60].

### Does male-male competition moderate the effects of past reproductive investment?

To test if male-male competition moderates the observed reproductive costs on later-life performance, we ran an additional set of analyses (Supporting Information) to compare a situations when two male rivals were present (this study) or absent [32]. In the current study, *ablated* and *non-ablated* males interacted freely with two rivals, while *naïve* males had visual access to two rivals. In the earlier study [32], focal males were only housed with a female. Data was only available to test for an effect of rivals after 8 weeks.

Male-male competition did not exacerbate the reproductive costs of pursuing females or releasing sperm (S4 and S5 Tables). For seven of the eight measured traits, there was no interaction between the reproductive treatment and the presence/absence of rivals. In the combined data, greater past mating effort led to slower growth, lower immunocompetence, and fewer mating attempts by *ablated* and *non-ablated* males (Tukey’s tests: versus *naïve* males: all p < .013) (S5 Table), but past sperm release did not have these effects (Tukey’s tests: *ablated* vs *non-ablated*, all p > .210). Likewise, past sperm release led to a significantly lower sperm count and sperm replenishment rate in the combined data (Tukey’s tests: *non-ablated* males vs *naïve* or *ablated* males: all p < .013), but past mating effort did not have these effects (Tukey’s tests: both p > .264; S5 Table). The sole exception was the time spent near females. In the absence of rivals, past reproductive investment did not affect the time spent near females [32]. In the presence of rivals, however, males with greater past mating effort (*ablated* or *non-ablated*) spent significantly less time near females than *naïve* males; while past sperm release had no effect (Fig 2B). This suggests that antagonistic interactions with rivals and pursuing females have cumulative costs on a male’s ability to pursue females, perhaps because they both involve locomotion that causes muscle damage.

In sum, there is robust evidence using the data from two independent studies that: (1) past mating effort reduces growth and lowers immunity; (2) past mating effort and past sperm release lower subsequent reproduction in ways that differ predictably among traits: ejaculation lowers future sperm production, while mating effort lowers future mating performance. Interestingly, antagonistic interactions with rivals did not moderate how past reproductive investment affected future ejaculate production. As noted earlier, this is consistent with the costs of repeated ejaculation being confined to germline tissue, which would not be directly affected by chasing or fighting rivals. The two studies were conducted at different times, albeit using the same population of fish and identical laboratory facilities, so caution is needed when comparing their results. Even so, it worth noting that males that had previously encountered rivals (i.e., this study) replenished their sperm significantly more rapidly and produced faster sperm (even for *naïve* males that only saw rivals) than males that did not encountered rivals [32] (S5 Table). This difference is consistent with males adjusting their ejaculate size to the perceived risk/intensity of sperm competition (meta-analysis: [13]).

## Conclusion

We ran an experiment to disentangle how pre- and post-copulatory investment affect male life history traits when males compete for mates and fertilization. Our key finding is that the long-term costs of mating effort and ejaculation are such that there is no detectable trade-offs between pre- and post-copulatory traits. This finding cautions against the generality of life-history models that assume such a trade-off to then predict how males optimize allocation between mate acquisition traits and sperm competitiveness according to, for example, the risk/intensity of sperm competition [20] or a male’s own competitiveness [19]. More generally, social and ecological factors that differentially affect the level of sperm and mating competition can lead to adaptive plastic shifts in reproductive investment into pre- and post-copulatory traits [5]. This could subsequently generate variation in the rate of senescence of different sexually or naturally selected traits that is not predicted if it is assumed that pre- and post-copulatory investment in sexual traits has equivalent life history effects.

## Materials and Methods

### Origin and maintenance of fish

In May 2020, mosquitofish (*G. holbrooki*) were collected from ponds in Canberra, Australia (35° 180 27” S 149° 070 27.9” E) (ACT Collection license FS20188) and housed in aquarium facilities at the Australian National University (permit A2021/04). Adults were introduced into single-sex stock 90*l* tanks (40-50 fish/aquarium). Juveniles were raised communally until their sex could be determined prior to maturation (an elongated anal fin for males; visible gravid spot for females). Males and females were then separated into single-sex 90*l* tanks to ensure virginity. All fish were maintained under 14:10 L:D photoperiod at 28 °C and fed twice daily with *Artemia* nauplii *ad libitum* and commercial fish flakes if in stock tanks, or only *Artemia* when in individual tanks. The experiment ran from June 2020 to January 2021.

### Experimental design

Male mosquitofish almost exclusively employ a coercive mating strategy: a male approaches a female from behind, swings his gonopodium (intromittent organ) forward, and insert the tip into her gonopore to transfer sperm [35]. We screened for sexually active males by randomly selecting a virgin male and placing him into a 4*l* tank with a wild-caught female for 5 minutes. Only males who chased the female and attempted to mate were included as focal males. We then placed the male in an ice-slurry for 10 seconds as anaesthesia before photographing him to measure his standard length (SL) and body depth (BD). Photographs were analysed using *ImageJ* [61]. Size-matched males (ANOVA, SL: *F* = 1.228, p = .296; BD: *F* = 2.369, p = .100) were haphazardly assigned to three treatments: (a) “Naïve”: a focal non-ablated virgin male, a wild-caught female and two wild-caught rival males were placed into three separate chambers of a 7*l* tank using mesh barriers. The focal male experienced only olfactory and visual cues from the female and the rivals without any physical contact, so he did not fully invest into mating behavior nor did he ejaculate (hence low/no need for sperm replenishment) (*n* = 56); (b) “Mating only” (‘ablated male’): a focal ablated virgin male (see below for ablation surgery) was placed with a female and two rival males. The focal male engaged in aggressive interactions with the rivals, and attempted to chase and mate the female, but he did not ejaculate (i.e., low/no need for sperm replenishment). He mainly allocated resources to mating effort (*n* = 54); (c) “Mating and ejaculation” (‘non-ablated male’): a focal non-ablated virgin male was housed with a female and two rivals. The focal male could fight with the rivals, chase the female and ejaculate, thereby investing into mating effort, ejaculation and sperm replenishment (*n* = 53).

Males from all three treatments paid the cost of baseline sperm production, however, “mating and ejaculation” *non-ablated* males had to replenish sperm more often due to repeated sperm release each time they mated. This, in turn, promotes a higher division rate in the germline, as well as any maintenance costs of spermatozoa [24]. Although we cannot exclude the possibility that *naïve* or “mating only” *ablated* males discard and/or reabsorb unused sperm, poecilid fish infrequently refresh stored sperm reserves [62]. As such, ejaculate investment by *naive* and *ablated* males should be far lower than that for *non-ablated* males because they ejaculated and replenished sperm more often. A comparison between *ablated* and *non-ablated* males is therefore a biologically meaningful estimate of the cost of repeated ejaculation during actual mating.

We anesthetized a focal male using iced water for 10 seconds, placed him on a glass slide and swung his gonopodium forward under a dissecting microscope. According to his treatment, we either ablated or sham-ablated the tip of his gonopodium with a scalpel blade (Diplomat Blades, Victoria, Australia). The removal of the tip prevents a male from receiving the cues that trigger ejaculation [31]. Males were then transferred into individual 7*l* tanks for a 3-day recovery period, after which we introduced a stimulus female and two wild-caught stock males (see above). Focal males were labelled using numbering so that we were blind to their treatment. Stimulus females and male rivals were rotated between tanks weekly to minimize effects of female familiarity, and variation in the focal males’ dominance status. To identify the focal male, all rival males were anaesthetized in ice slurry and marked with an elastomer tag (Northwest Marine Technology, Shaw Island, WA) injected subcutaneous below the dorsal fin.

The treatments were maintained for 16 weeks. At the end of weeks 8 and 16, somatic and reproductive traits were measured (see below). Ablated males did not re-grow their gonopodium tip (personal observation) nor did gonopodium length increase with age (LM, age at measurement: *F* = 1.582, p = .209, with week 0 gonopodium length as a covariate: *F* = 1331.286, p < .001).

### Preliminary test

Virgin males were randomly assigned to be *ablated* or *non-ablated* (*n* = 70 per treatment). After a 3-day recovery (Day 0), we introduced a focal male (ablated or non-ablated), an intact rival male (21.06 ± 0.14 mm SL; *n* = 140) and a wild-caught female (27.98 ± 0.20 mm SL; *n* = 140) into a 7*l* tank. The three individuals were separated by two mesh barriers for 10 minutes. We then raised the barriers and recorded the following behaviors of the focal male for 20 minutes: (a) number of mating attempts, (b) time spent within 1 SL of the female, (c) number of nips and rapid approaches to the rival male. After the trial, each focal male was housed in a 7*l* tank with a female and two rivals for 7 days to allow for sexual interaction (with or without ejaculation depending on his treatment). On Day 7, we re-ran the behavioral trial using a new rival (21.13 ± 0.13 mm SL; *n* = 130) and a novel female (28.12 ± 0.19 mm SL; *n* = 130) to test for any differences in mating performance. Rivals were marked with an elastomer tag.

### Body growth

Males were photographed under anaesthesia to measure their SL and BD after 8 and 16 weeks in the treatments. Growth was determined by their current SL and BD controlling for their previous size.

### Immune response

Cell-mediate immunity was tested using a phytohaemagglutinin injection assay [28]. We used a pressure-sensitive spessimeter (Mitutoyo 547-301, accuracy: 0.01 mm) to measure the body thickness at the posterior end of the dorsal fin of males under anaesthesia. We made five measures per male and used the average value in our analyses. We then injected 0.01mg phytohaemagglutinin (dissolved in 10μl PBS) into the left side of the fish at the point where thickness was quantified. After 24 hours, we re-measured the male and calculated the difference between the pre- and post-injection values as our measure of his immune response.

### Mating performance

After 8 and 16 weeks in their assigned treatments, we examined the effect of reproductive investment on the focal males’ mating behavior by recording: (a) number of nips and rapid approaches, (b) time spent near the female, and (c) number of mating attempts over 20 minutes following the same protocol used in the *Preliminary test*.

### Sperm traits

We tested for any differences in sperm traits due to the reproductive treatment after 8 and 16 weeks. To control for variation in sperm age, we first emptied all sperm reserves of focal males (Day 0) and isolated them in separate 7*l* tanks for 5 days to produce new sperm bundles [36]. On Day 5, we put an anaesthetized male on a glass slide covered by 1% polyvinyl alcohol solution, swung the gonopodium forward under a dissecting microscope and gently pressed on his abdomen to eject sperm bundles. We collected the sperm into a known volume (200–1200μl depending on ejaculate size to optimize the count measurement) of extender medium (207 mM NaCl, 5.4 mM KCl, 1.3 mM CaCl_2_, 0.49 mM MgCl_2_, 0.41 mM MgSO_4_, 10 mM Tris (Cl); pH 7.5). After vortexing the solution, we placed 3μl on a 20-micron capillary slide (Leja) under 100× magnification. Sperm number was calculated using the automated program CEROS Sperm Tracker (Hamilton Thorne Research, Beverly, MA, USA) to minimize the risk of observer bias. After 24 hours (Day 6), we again stripped the male and counted the number of sperm as a measure of his daily rate of sperm replenishment. We determined the total sperm count and number of replenished sperm using the mean value of five randomly selected subsamples [29] (repeatability: *r* ± SE = 0.924 ± 0.005, p < .001, *n* = 595 male-days).

When testing sperm count, we also collected two samples per male to determine sperm velocity [29]. For each sample, we pipetted a 3μl solution of 3 sperm bundles and extender medium into the centre of a cell in a 12-cell multi-test slide (MP Biomedicals, USA) coated with 1% polyvinyl alcohol solution. The sample was activated with 3μl of 125 mM KCl and 2 mg/ml bovine serum albumin for 30 seconds and covered with a coverslip. We recorded the velocity for 81.94 ± 2.92 *SE* sperm tracks per ejaculate and measured (a) curvilinear velocity (VCL): the actual velocity along the trajectory, (b) average path velocity (VAP): the average velocity over a smoothed cell path and (c) straight-line velocity (VSL) using the CEROS Sperm Tracker. Given that all three measures were highly correlated (VCL-VAP: *r* = 0.998; VCL-VSL: *r* = 0.999; VAP-VSL: *r* = 0.999; *n* = 300), we report the actual velocity (VCL).

### Statistical analysis

Our analysis plan was pre-registered online (osf.io/swdv7).

#### Preliminary test

Given the sample distribution of each trait, separate generalized linear mixed models (GLMMs) with quasi-Poisson error were used for: (a) number of mating attempts and (b) number of nips and approaches to rival, while (c) time spent near the female was analyzed using a linear mixed model (LMM). We considered the state of the gonopodium (ablated, non-ablated), test day (Day 0, Day 7) and standardized SL of focal male (mean = 0, SD = 1) as fixed factors. We included all two-way interactions (gonopodium state*day, day*focal male size, gonopodium state*focal male size) in our initial models. Male identity was treated as a random factor to account for repeated measurements from each male. We included the standardized size difference between the focal and the new rival male in behavioral test as a covariate because relatively larger males tended to be more aggressive [63]. If gonopodium state and test day showed a significant interaction, we ran separate generalized linear models (GLMs) with negative binomial error for each day to test for differences in behavior between *ablated* and *non-ablated* males.

#### Main experiment

We ran separate LMMs for: (a) standard length, (b) body depth, (c) immune response, (d) time spent near female, (e) total sperm count (log-transformed), (f) sperm replenishment rate (log-transformed) and (g) sperm velocity. We ran separate GLMMs with quasi-Poisson error for: (h) number of mating attempts (accounting for zero-inflation) and (i) number of nips and rapid approaches. Reproductive treatment (naïve, ablated, non-ablated) and age at testing (week 8, week 16) were considered as fixed factors, and we included their two-way interaction in initial models. If there was a significant interaction, we ran separate general linear models (LMs) for weeks 8 and 16 to test if the costs of reproduction changed with age at testing.

We included the standardized body size of focal males before the first and second half of the treatment period as a covariate to account for any effect of pre-treatment body size on subsequent mating performance and sperm traits (S1 Fig). We included male identity as a random factor for most traits, expect for (b) body depth and (g) sperm velocity, because the variance attributable to male identity was nearly zero in these mixed models. We instead ran simple LMs for these two traits. Finally, we included the standardized size difference between the focal and rival male as a covariate for the analyses of mating performance.

In all models, non-significant interactions were removed to interpret main effects [64]. We also confirmed that removal of non-significant interactions did not significantly lower the model fit (S6 Table). Tukey’s post-hoc tests (*emmeans* package) were run to test for pairwise difference between treatments. We confirmed that the analyses of LMs and LMMs fulfilled model assumptions (i.e., homoscedascity and normally distributed residuals) using Q-Q plots. We conducted dispersion test (*DHARMa* package) to ensure the data variance fulfilled the model assumption in GLMs and GLMMs. Results are presented as mean ± *SE* and the significance level was set at alpha = 0.05 (two-tailed) using the *Anova* function (type III Wald *chi-square* tests for GLMs and GLMMs; *F*-tests for LMs and LMMs) in the *car* package of R v4.0.5 [65] using Rstudio v1.3.1093.

## Acknowledgements

We thank Ben Durant, Michelle Stephens, Upama Aich, Lauren Harrison and Ivan Vinogradov for help with fish maintenance.

## Ethic statement

The project (including fish maintenance, ablation surgery, trait measurement) received approval from the Australian National University’s Animal Ethics Committee (Ethics Protocol: A2021/04).

## Supporting information

Supporting information has been uploaded separately as an individual file.

**S1 Fig. Relationship between initial body length and male reproductive traits.** (A) number of mating attempts; (B) total sperm count; (C) rate of sperm replenishment; (D) sperm velocity.

**S1 Table**. Statistical outputs of initial models including all interaction terms for the surgery effect (i.e., gonopodium state) on mating behaviors in the presence of male rivals. The bold font highlights significance at the 0.05 level.

**S2 Table**. Statistical outputs for interactions and main effects of reproductive treatment and age at testing on somatic traits. Given a significant interaction, the main effects were tested separately at weeks 8 and 16. The bold font highlights significance at the 0.05 level.

**S3 Table**. Statistical outputs of initial models including the interaction between reproductive treatment and age at testing on reproductive traits. The bold font highlights significance at the 0.05 level. Final models excluding the non-significant interaction were provided in Table 1 of the main text.

**S4 Table**. Statistical outputs of initial models including the interaction between reproductive treatment and the presence of rivals on somatic and reproductive traits. The bold font highlights significance at the 0.05 level. The model of *time spent near female* and final models excluding the non-significant interaction were presented in S5 Table.

**S5 Table.** Effects of reproductive treatment and the presence of rivals (i.e., levels of male-male competition) on trait expression after 8 weeks. Main effects are obtained from final models excluding non-significant interactions, except for *time spent near female* which had a significant interaction. Statistical outputs of initial models with the interaction are provided in S4 Table.

**S6 Table.** Comparison of the best fit of the models.

